# Hundreds of Putative Non-Coding Cis-Regulatory Drivers in Chronic Lymphocytic Leukaemia and Skin Cancer

**DOI:** 10.1101/174219

**Authors:** Halit Ongen, Olivier Delaneau, Michael W. Stevens, Cedric Howald, Emmanouil T. Dermitzakis

## Abstract

Perturbations of the coding genome and their role in cancer development have been studied extensively. However, the non-coding genome’s contribution in cancer is poorly understood (*1*), not only because it is difficult to define the non-coding regulatory regions and the genes they regulate, but also because there is limited power owing to the regulatory regions’ small size. In this study, we try to resolve this issue by defining modules of coordinated non-coding regulatory regions of genes (Cis Regulatory Domains or CRDs). To do so, we use the correlation between histone modifications, assayed by ChIP-seq, in population samples of immortalized B-cells and skin fibroblasts. We screen for CRDs that accumulate an excess of somatic mutations in chronic lymphocytic leukaemia (CLL) and skin cancer, which affect these cell types, after accounting for somatic mutational patterns and biases. At 5% FDR, we find 90 CRDs with significant excess somatic of mutations in CLL, 60 of which regulate 126 genes, and in skin cancer 59 significant CRDs, 25 of which regulate 37 genes. The genes these CRDs regulate include ones already implicated in tumorigenesis, and are enriched in pathways already implicated in the respective cancers, like the B-cell receptor signalling pathway in CLL and the TGFβ signalling pathway in skin cancer. We discover that the somatic mutations in the significant CRDs of CLL are hitting bases more likely to be functional than the mutations in non-significant CRDs. Moreover, in both cancers, mutational signatures observed in the regulatory regions of significant CRDs deviate significantly from their null sequences. Both results indicate selection acting on CRDs during tumorigenesis. Finally, we find that the transcription factor biding sites that are disturbed by the somatic mutations in significant CRDs are enriched for factors known to be involved in cancer development. We are describing a new powerful approach to discover non-coding regions involved in tumorigenesis in CLL and skin cancer and this approach could be generalized to other cancers.

## INTRODUCTION

The Cancer Genome Atlas Consortium (TCGA) (*2*) and International Cancer Genome Consortium (ICGC) (*3*) have so far been tremendous resources to generate and characterize genomics data from a variety of human cancer types. By investigating somatic mutations in the protein coding parts of the genome, they have identified many genes as putative driver genes across multiple cancers (*4*), as well as many known and novel putative drivers in specific cancers (*5–11*). However, with the whole genome sequencing (WGS) data generated by these projects, it became evident that most of the somatic mutations observed lie in the non-coding portion of the human genome (*1*). One important question that follows from this observation is whether these non-coding somatic mutations are involved in tumorigenesis, or whether they are simply passenger mutations that are not contributing to tumorigenesis and are therefore not under positive selection in cancer development.

Unlike the coding genome’s impact on tumorigenesis, the contribution of the non-coding genome to cancer development is less well understood, since interpreting the non-coding genome remains challenging (*1*). A few studies identified non-coding regulatory mutations are involved in tumorigenesis such as the *TERT* promoter in melanoma (*12, 13*) and in other cancers (*14*), and recurrent non-coding mutations in chronic lymphocytic leukaemia (CLL) (*15*) and breast cancer (*16*). However, these studies have each identified a small number of regulatory regions mainly due to power issues, as regulatory elements are individually studied and tend to be small in size. Recently it has become evident the non-coding mutations play an important role in tumorigenesis (*17*). We have also previously shown that non-coding germline variation is likely involved in tumorigenesis, and identified genes with putative somatic regulatory drivers in colorectal cancer, using perturbations in allele specific expression during tumour development (*18*). However, directly finding non-coding regulatory regions that accumulate an excess of somatic mutations using WGS data is an unsolved problem due to the difficulty in defining the bounds of the non-coding regulatory regions, assessing which genes these regulate, identifying the null genome to compare to, and failure to accumulate signal over multiple regulatory regions thus decreasing power in detecting these regions. In this study, we address this problem by identifying sets of coordinated non-coding regulatory regions, called cis regulatory domains (CRDs), based on the correlation amongst three histone modifications, and by finding which genes these CRDs regulate. We then test whether these CRDs accumulate an excess of somatic mutations by controlling for differential somatic mutation rate across the genome and between open and closed chromatin regions.

## RESULTS

### Identification of cis regulatory domains

In order to define likely regulatory non-coding regions of the genome we have used data generated by the SysGenetiX (SGX) project (*19*), which comprises genotypes, H3K27ac, H3K4me1, H3K4me3 histone modifications assayed by ChIP-seq, and transcriptomes assayed by RNA-seq from 317 lymphoblastoid cell line (LCL, immortalized B-cells) and 78 skin fibroblast cell lines. With this dataset, we can use the correlation between the chromatin marks in the population to identify likely regulatory regions of the genome in a cell type specific manner. To achieve this, we computed all the pairwise correlations between nearby chromatin peaks, which were grouped into a tree using hierarchical clustering. We defined cis regulatory domains (CDRs) as nodes on the tree where the mean correlation of the leaves of the node is double the background correlation and find 21607 and 18261 CRDs in LCLs and skin fibroblasts, respectively (these counts are higher than the original SGX study, since we are also considering CRDs comprised of overlapping peaks). Furthermore, using the resulting CRD and gene expression quantifications we were able to discover 15161 and 3652 nearby genes (FDR = 5%, 1 Mb cis window) that the CRDs regulate in LCLs and fibroblasts, respectively (**Supplementary methods & Supplementary figure 1**). By finding CRDs and genes that covary in the population, we can estimate the non-coding regulatory regions of genes, which will not only enable identification of non-coding regulatory regions with excess somatic mutations but also link these regions to specific genes.

### Identifying CRDs with excess somatic mutations accounting for somatic mutation biases

Next, we devised a method to detect excess somatic mutations in CRDs, which would signify positive selection in tumorigenesis and hence indicate driving potential. To do so we first identified the regulatory regions of the CRDs from the SGX project (*19*) that do not overlap with known exons. Subsequently, we created a set of regions that did not overlap with any of the ChIP-seq peaks in SGX or with known exons, which make up non-exonic non-regulatory null regions that we call spacers. In order to account for differential somatic mutation rates observed across the genome (*20–23*), we took the spacers in-between or flanking the regulatory regions of a CRD as the null for a given CRD. Activation induced deaminase (AID) is known to cause hypermutation in lymphomas (*24, 25*) and excess mutations observed in these regions may be due to hypermutation rather than positive selection. To account for this in our model, we categorized the somatic mutations seen in CLL to canonical and non-canonical AID signatures and measured the AID mutation rate across the genome. We then segmented the genome into regions with similar AID mutation rates and scaled the AID mutations observed in these regions by the ratio of non-AID mutation rate to AID mutation rate (**Supplementary methods, Supplementary figure 12 & 13**). This approach allowed us to penalize individual AID mutations in hypermutated regions of the genome, and effectively normalize the AID mutation rate across the genome. Different cancers differ in their mutational signatures, i.e. in the types of somatic mutations they accumulate (*26*). In order to incorporate this into our method, we considered the local context of the somatic mutations observed by taking into account the immediate 5’ and 3’ reference bases (trinucleotide context) flanking the somatic mutation position. These trinucleotide contexts were summarized on the pyrimidine bases of the observed somatic mutation. Thus, we counted how many of the 32 possible context triplets are present in the spacer regions of a CRD and how many of them harbour a somatic mutation, to calculate an expected mutation rate for each of the 32 classes. We then counted the number of times these triplets are observed in the regulatory regions of a CRD and using the expected mutation rates for each of the 32 classes, we found the total number of expected mutations. Subsequently we counted the observed number of mutations in the regulatory regions and using a one-tailed Poisson test calculated a nominal p-value for the excess of somatic mutations in a CRD. Lastly, we wanted to account for differential mutation rates and other unknown technical biases between open (regulatory regions) and closed chromatin (spacers) (*27*). Thus, we devised a permutation scheme, in which for a triplet context in the regulatory region of a CRD we select a random position with the same context in a random CRD and its corresponding downstream spacer and ask if these positions harbour somatic mutations. We did this for all of the regulatory bases of a CRD, to generate a pseudo CRD, which conserves the differential mutation rates in open vs. closed chromatin, but breaks the clustering of the somatic mutations. Then, we iterate this process 100000 times for each of the CRDs and at each iteration assessed the significance of the observed nominal p-value relative to the permuted p-values to come up with our adjusted p-value (**Supplementary methods, Supplementary figure 2 & 3**). Please see supplementary methods section 3.1, 3.2, and 3.3 for the details of these steps and section 3.4 for how we accounted for all the known biases. Using this methodology, we controlled for the differential mutation rate across the genome, hypermutation observed in CLL, the local context of the types of somatic mutations, and the different rates of somatic mutation inferences between open and closed chromatin, therefore resulting in a robust estimate of the significance of excess somatic mutations in CRDs.

### CRDs with excess somatic mutations

We aimed to find CRDs with significant excess of somatic mutations with the aforementioned methodology using publicly available somatic mutation calls. To this end, we acquired somatic mutation calls from WGS data for CLL (*15*) and through the International Cancer Genome Consortium data portal for skin cancer (project codes: MELA-AU and SKCA-BR) (*28*). These cancers were chosen because they affect the cell types analysed in the SGX project. In CLL we found 90 CRDs that accumulate significantly more somatic mutations (FDR = 5%), 60 of which regulate 126 genes, and in skin cancer we found 59 significant CRDs (FDR = 5%), 25 of which regulate 37 genes (**Figure 1**, **Supplementary tables 1 & 2**). Due to limitations in power, not all modules are assigned to genes, which is more apparent in the smaller sample sized skin fibroblasts.

**Figure 1.**
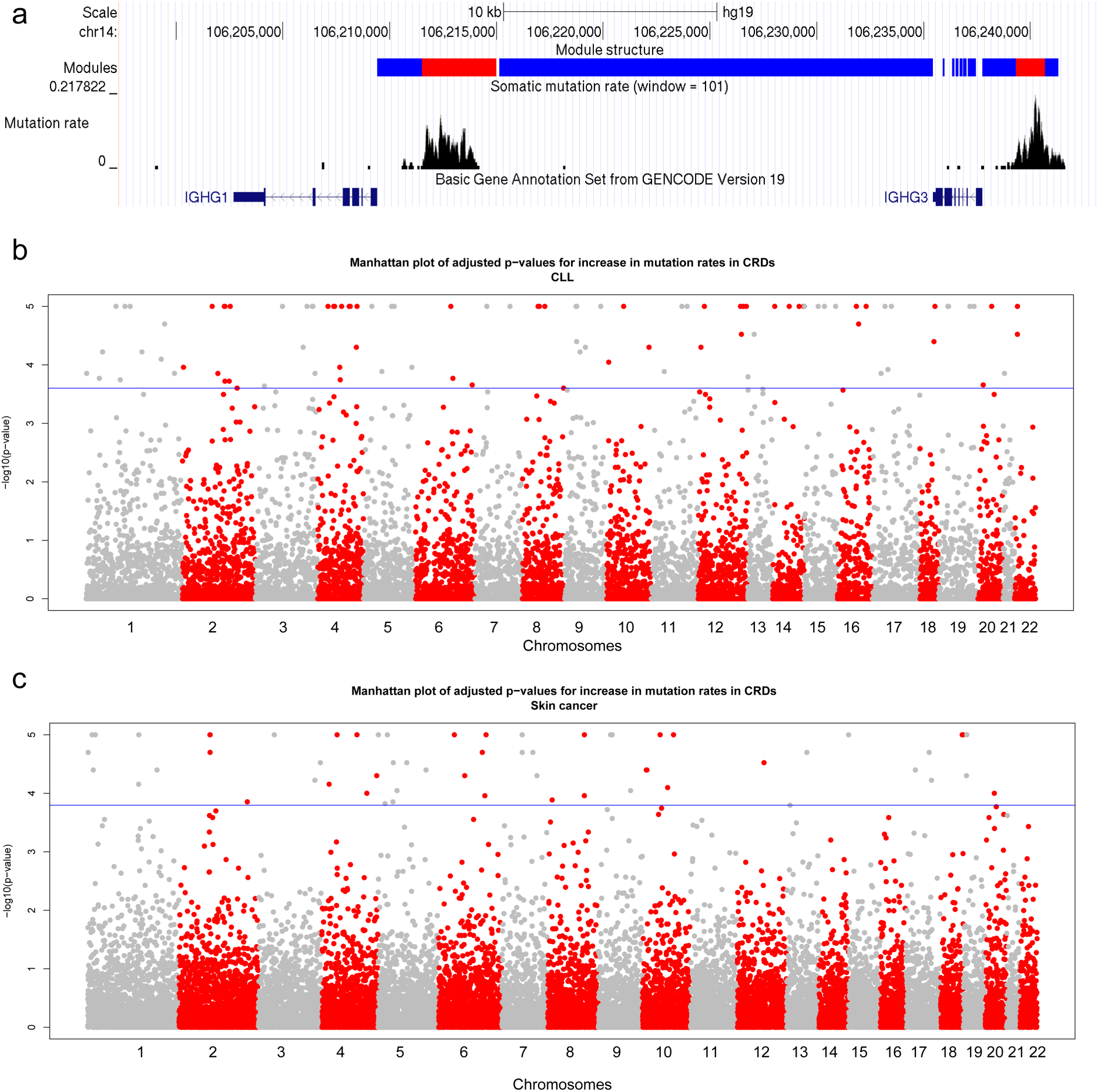
– (a) An example of a CRD that has significantly excess somatic mutations in CLL, which regulates the expression of the *IGHG3* gene. The blue red boxes are the positions of the CHiP-seq peaks of the CRD, i.e. the regulatory regions of this CRD, and the blue boxes are the spacers. The mutation rate averaged over a sliding window of 101 bp is shown as black bars. The gene structure in the region is represented as dark blue boxes. Plot is generated using the UCSC genome browser. Manhattan plot of the permuted p-values for increase in mutation rates in CRDs for (b) CLL and (c) skin cancer. The blue line indicates the 5% FDR threshold. Alternating chromosomes are coloured differently.

One possible technical issue that might confound our analysis is coverage in WGS. If the regulatory regions of the significant CRDs had systematically higher coverage than their spacers, then this would cause more somatic mutations to be called in regulatory regions. To assess this, we compared the number of reads overlapping the somatic mutation calls in the regulatory regions of the significant CRD to the number of reads of around somatic mutations in their spacers. In both cancer types, there was no significant increase in coverage of somatic mutations in regulatory region compared to their spacers (**Supplementary figure 9**). In fact, the coverage in the regulatory regions was lower than in the spacers in both cancers (median 21 reads in regulatory vs. 31 in spacer in CLL and 60 vs. 61 in skin cancer). We also investigated whether inclusion of different spacers, e.g. including or excluding flanking spacers, might influence our results.; thus we reran the analysis without the flanking regions. We observed that the p-values were significantly positively correlated between the two analyses (rho = 0.825, p < 2e-16 in CLL and rho = 0.404, p = 0.008 in skin cancer). Skin cancer showed a higher difference between the two analyses but in almost all cases inclusion of the flanking spacers made the p-value less significant (**Supplementary figure 10**). These results indicate that our permutation scheme accounts for any heterogeneity of coverage and variant call confidence and that our results hold with different ranges of spacer sequence inclusion.

We examined the per sample somatic mutation rates of the significant CRDs, and observed that similar to the protein coding drivers there are CRDs that are highly mutated in most samples, and others exhibiting high mutation rates in just few samples (**Figure 2**). This suggests that the patterns of selection in non-coding cis-regulatory regions are similar to protein coding sequences with some regulatory regions probably important for tumorigenesis in most or all cancers while others being more specific to one or a few tumors.

**Figure 2.**
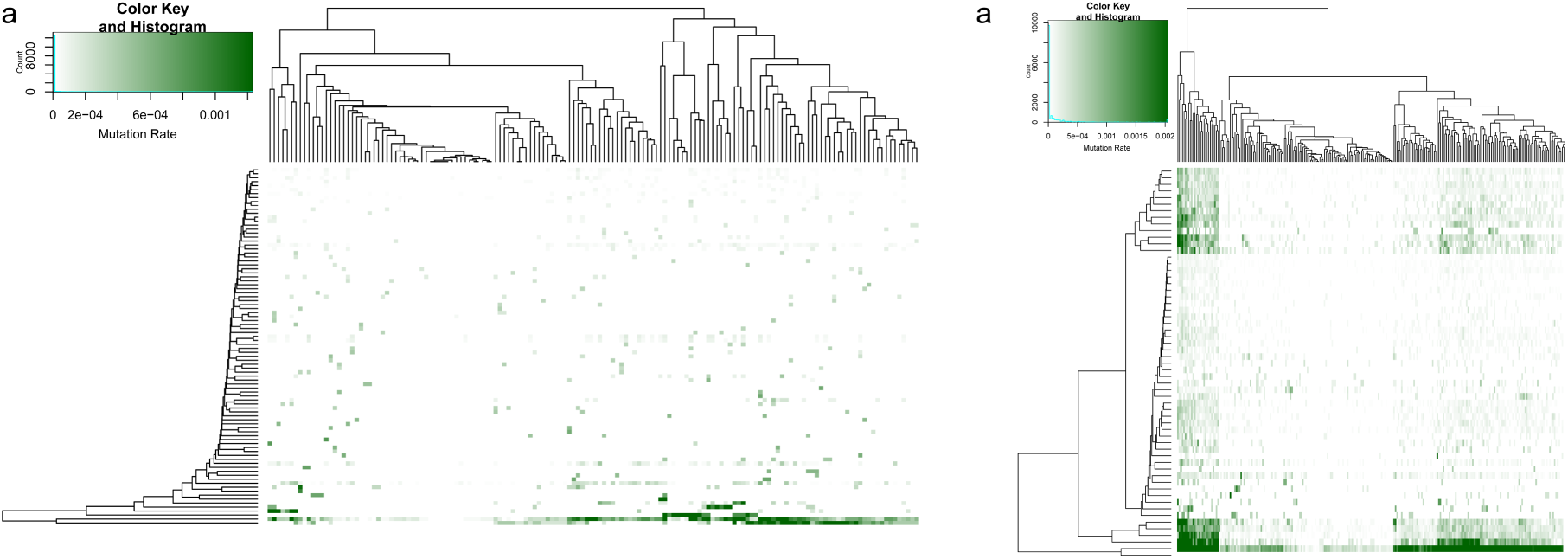
– The clustering of samples based on the somatic mutation rates of the significant CRDs in CLL (a) and in skin cancer (b). The rows are the CRDs and the columns represent samples. The hierarchical clustering of samples and CRDs are shown as a dendrograms. The clustering is conducted with hierarchical clustering using complete linkage method on the Euclidian distances calculated from the mutation rates.

Among the genes identified in CLL are *BIRC3* (**Supplementary figure 4**), a putative driver previously identified due to excess protein altering mutations (*29*), *BCR*, which is one of the breakpoints of the Philadelphia chromosome first identified in chronic myeloid leukaemia (*30*), and *LPP*, where the CRD contains a variant that confers predisposition to CLL (*31*). Overall the genes identified in CLL are enriched for B-cell receptor signalling pathway, one of the major pathways involved in CLL development (*32*), and for pathways involved in the tumorigenesis of other types of cancers (**Supplementary table 3**). Nine of the significant CLL CRDs are in the hypermutated *IGHV* locus; however not only we account for this hypermutation rate in our analysis but also, we show below using the functional assessment of CLL somatic mutations, that these are unlikely to be false positives. In skin cancer we find CRDs for genes such as *LRRC37A3* (**Supplementary figure 5**) and *TMEM200A*, which have been previously identified as putative drivers in melanoma due to an excess of protein altering mutations (*6*). Collectively these genes are enriched for the TGFβ signalling pathway (**Supplementary table 4**), an important pathway in skin cancer development (*33*). We show that by using our methodology we can identify putative non-coding regulatory drivers for genes that are known to be involved in cancer development as well many novel ones.

### Comparison of significant CRDs vs. non-significant ones in both cancer types

Several patterns differ between the two cancer types. First, because of sample size differences between the cell types of the SGX project where skin fibroblasts have about four times smaller sample size, there is differential power in both discovering CRDs and assigning them to genes. Furthermore, for CLL we are directly assessing CRDs from the cell type that is giving rise to the cancer, whereas in skin cancer we are using the skin fibroblasts as the closest proxy available to the underlying cell type involved in tumorigenesis. Also, the underlying somatic mutation rate in skin cancer is significantly higher than in CLL. All these reasons individually and combined result in differences in our findings which we tried to highlight by comparing the significant CRDs to the non-significant ones in both cancer types.

We observed that in CLL, the average correlation between the chromatin peaks in significant CRDs is significantly higher than the correlation among the peaks in non-significant CRDs (**Supplementary Figure 11a**), indicating that the CRDs identified as cancer drivers indeed tend to have strong coordinated behaviour and therefore the accumulation of likely jointly contributing mutations to tumorigenesis is expected. In CLL the significant CRDs are significantly shorter, contain fewer chromatin peaks, regulate the expression of fewer genes that are further away from the CRD than in non-significant CRDs, highlighting again the discovery of strongly coordinated CRDs (**Supplementary Figure 11b, c, d, e, f**). Conversely, in skin cancer significant CRDs are longer, contain more chromatin peaks, regulate the expression of fewer genes that nearer to the CRD than in non-significant CRDs. The main reasons for this are on the one hand the smaller sample size in fibroblasts, which leads to fewer gene assignments and on the other hand, the increased mutation rate in skin cancer, which leads to signal dilution.

### Functional assessment of somatic mutations in the significant CRDs

We wanted to assess whether the somatic mutations found in the significant CRDs are perturbing likely functional nucleotide bases. In order to do so we acquired the probabilities for purifying selection acting on the bases of the human genome as calculated by the LINSIGHT method (*34*). Subsequently we compared the purifying selection acting on the positions of the somatic mutations in the significant CRDs vs. the positions in the non-significant CRDs (**Supplementary methods**). We found that, in CLL, somatic mutations in significant CRDs are hitting bases that are significantly (Mann-Whitney p < 2.2e-16) more likely to be under purifying selection than the mutated bases in non-significant CRDs, but this is not the case for skin cancer (**Figure 3**). We further investigated functional impact of the somatic mutations in the 9 significant CRDs overlapping the hypermutated *IGHV* locus in CLL (*29*). We found that the somatic mutations in the regulatory regions of these 9 CRDs are also hitting bases that are significantly more likely to be functional than both the mutated bases in non-significant CRDs (Mann-Whitney p < 2.2e-16) and the non-significant CRDs in the *IGHV* locus (Mann-Whitney p = 4.96e-6, **Supplementary figure 6**). The CLL result is indicative of selection in tumorigenesis, even in the hypermutated region, and the result for skin cancer, which on average has ~29 times more somatic mutations than CLL (Mann-Whitney p < 2.2e-16, **Supplementary figure 7**), is most likely due to the high number of somatic mutations diluting the relevant functional mutations.

**Figure 3.**
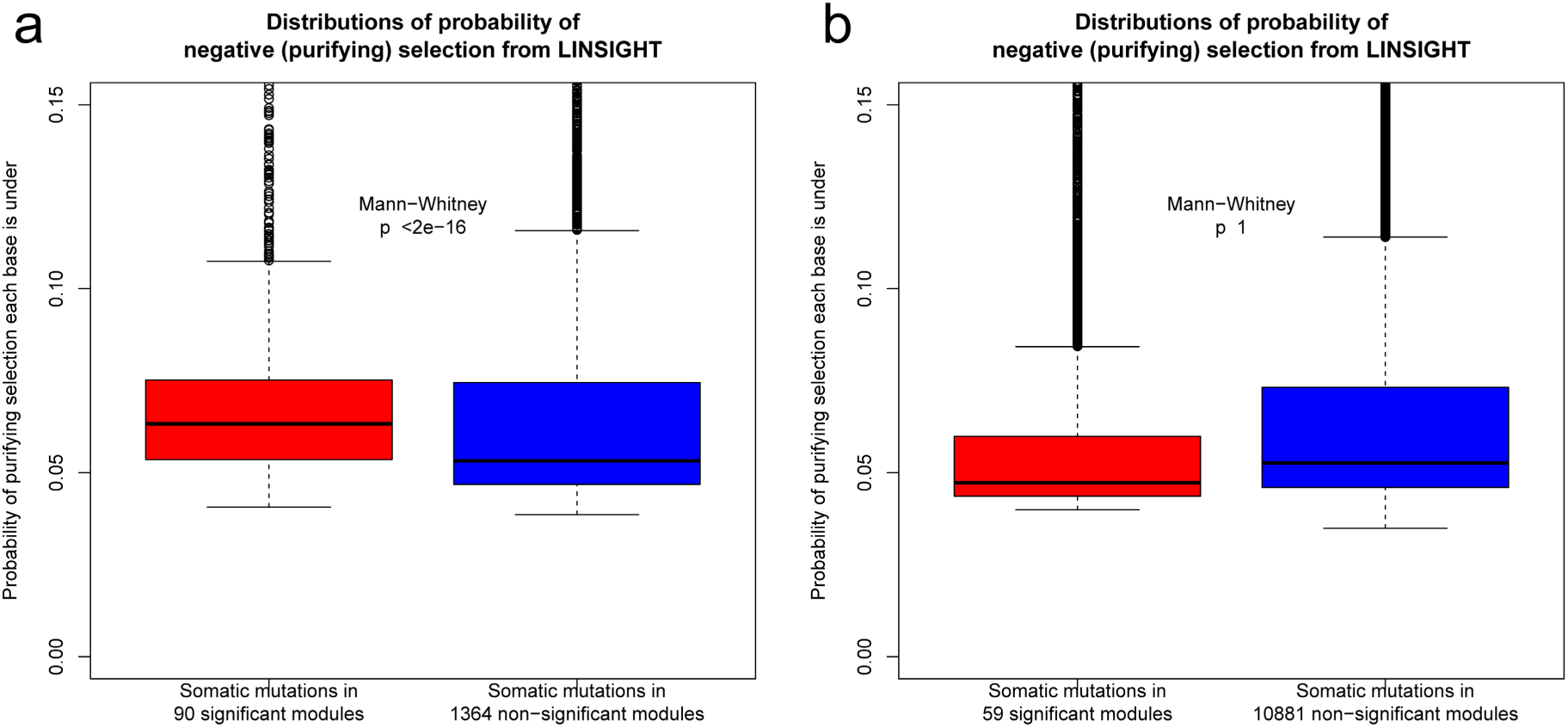
– The distributions of estimates of purifying selecting as calculated by LINSIGHT acting on somatic mutations in CRDs with significant excess of somatic mutations vs. ones that are not-significant in (a) CLL and (b) skin cancer. Boxplots are truncated at 0.15 probability of purifying selection. In CLL there is a significant increase in the purifying selecting acting on mutations in significant CRDs vs. non-significant ones, indicating that functional bases are being impacted in significant CRDs more so than in non-significant ones, suggesting selecting in tumorigenesis. This is not observed in skin cancer, likely due to very high number of somatic mutation seen skin cancer, diluting the signal.

To further investigate the selective pressure acting on the significant CRDs during tumorigenesis, we compared the types of somatic mutations observed between the regulatory regions and spacers in CRDs. To this end, we classified the single nucleotide somatic mutations into 96 classes representing six possible substitutions of the pyrimidine of the mutated Watson–Crick base pair with incorporating information on the immediate flanking bases (**Supplementary methods**) (*26*). If there is selective pressure on the significant CRDs, then we would expect the mutational signatures to differ between their regulatory regions and spacers, which we expect not to be under selection. In both cancer types we observed that there were significant differences in the mutation signatures in regulatory vs. spacer regions in significant CRDs (**Supplementary figure 8**). Furthermore, in skin cancer there was a significant decrease (Fisher’s exact test, OR= 0.83, p < 2.32e-14) in the C to T mutations, which are due to damage by U.V. light, in the regulatory regions vs. spacers of the significant CRDs. We also observed the same type of significant differences in the mutational signatures between the regulatory regions of the significant CRDs vs. the regulatory regions of the non-significant CRDs (**Figure 4**). These results signify that during tumorigenesis there is selection acting on the CRDs we identify independently of the underlying mutation rate, verifying that the signal of excess somatic mutations is not due to hypermutation, thus corroborating their driver potential.

**Figure 4.**
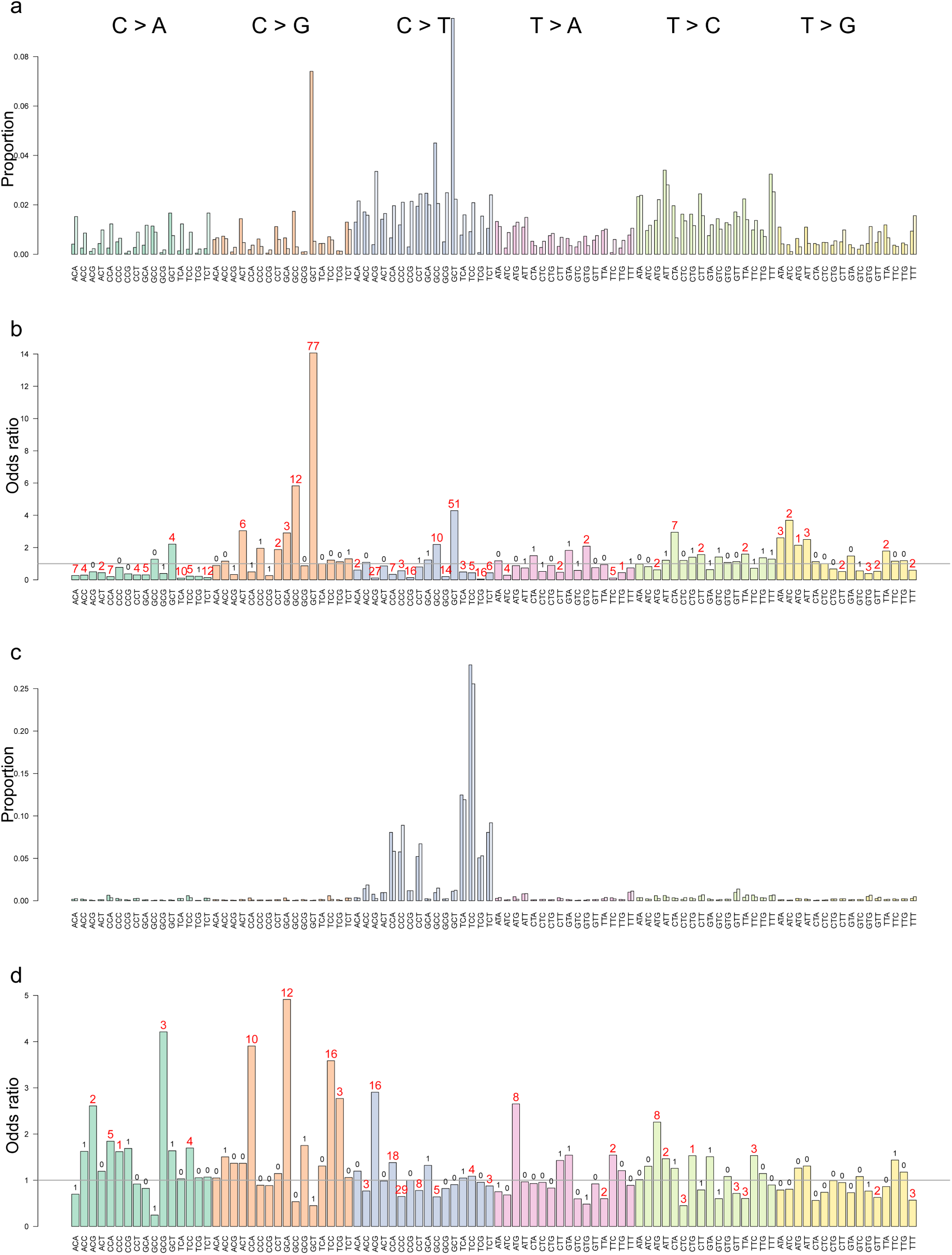
– The mutational signatures observed in the regulatory regions of the significant vs. non-significant CRDs. Bars are coloured by the nucleotide change observed and stratified by the local context. In (a, CLL) and (c, Skin cancer) the proportion of each mutational signature in the regulatory regions of significant CRDs are presented as darker shades whereas the non-significant CRDs are shown as lighter shades. In (b) and (d) odds ratio, proportion in significant CRDs’ regulatory regions over the proportion in non-significant CRDs’ regulatory regions, of a given mutational signature is plotted for CLL and skin cancer, respectively. The numbers above the bars are the Benjamini-Hochberg corrected –log10(p-value) of the observed enrichment or depletion where the significant classes are shown in red.

Lastly, we wanted to examine whether there are specific transcription factor binding sites (TFBSs) perturbed by the somatic mutations in the regulatory regions of the significant CRDs. Thus, we used sequences from regions flanking the somatic mutations by 15 bp (i.e 31 bp in total) in the regulatory regions of significant and non-significant CRDs. Then using the HOMER software package (*35*) we searched for TFBSs enriched around somatic mutations in significant CRDs compared to the non-significant CRDs (**Supplementary methods**). We found that in both CLL and skin cancer, transcription factors known to be involved in tumorigenesis are being perturbed, e.g. in CLL p53 (*36*), NFKB (*37*), E2A (*38, 39*), SCL (*40*), and in skin cancer CEBP (*41, 42*)(**Figure 5**). Therefore, with this analysis we identify changes in transcription factor binding sites as a likely mechanistic cause for the driver potential exhibited by these CRDs.

**Figure 5.**
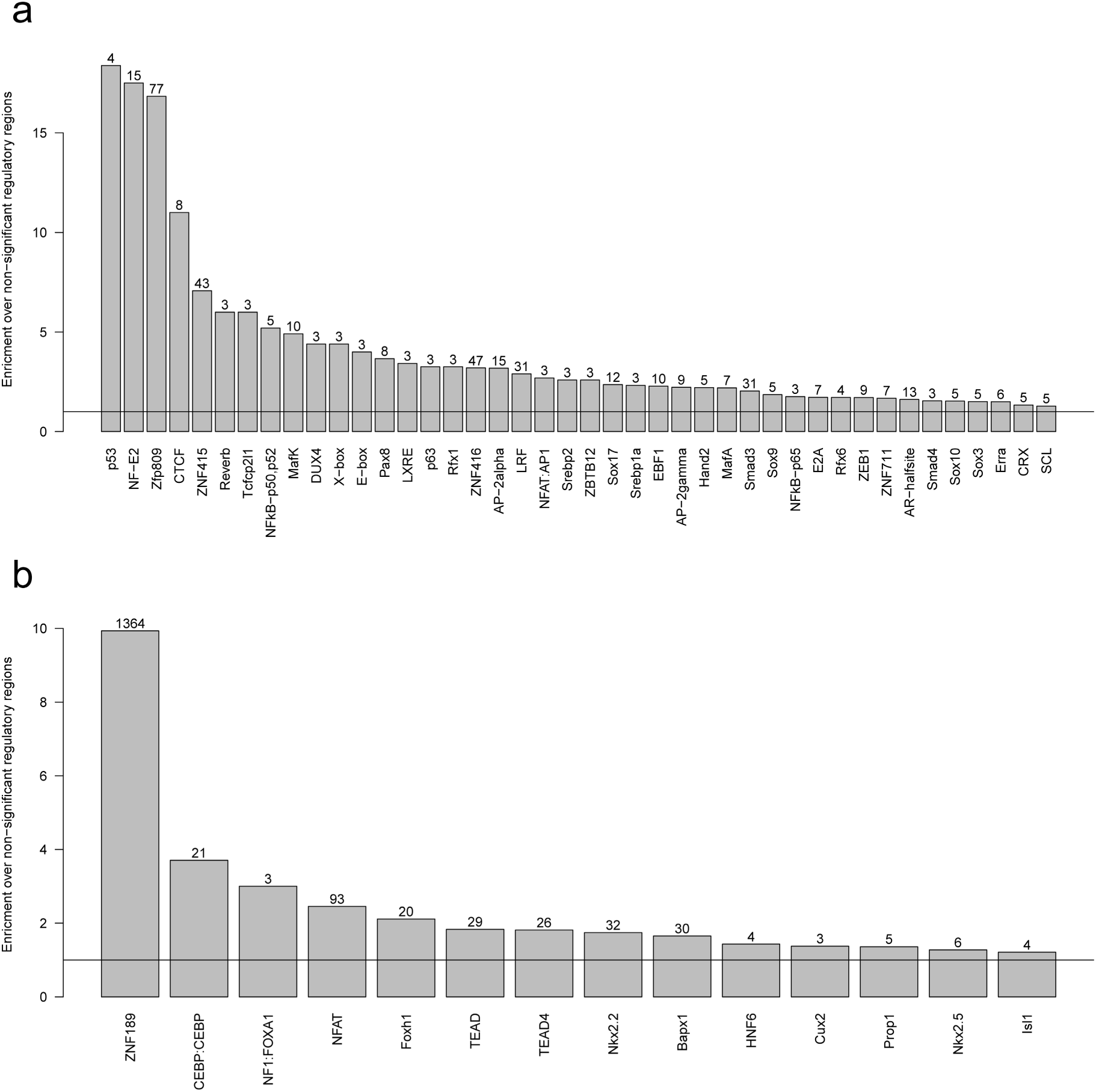
– The transcription factors that are significantly enriched around somatic mutations in significant CRDs vs. the ones in non-significant CRDs for (a) CLL and (b) skin cancer. The horizontal black line indicate the base line, and the numbers above the bars are the Benjamini-Hochberg corrected –log10(p-value) of the observed enrichment.

## DISCUSSION

In this study, we describe a methodology to identify non-coding cis-regulatory drivers in cancer in an unbiased and genome-wide manner using clusters of coordinated regulatory elements to increase power. We show that this method can find putative non-coding drivers in CLL and skin cancer, cancers affecting cell types analysed in the SGX project. We confirm our and other’s previous findings that non-coding drivers are important players in tumorigenesis (*17, 18*) and that the number of genes involved is much higher than currently known for protein coding drivers. As datasets like the SGX project or similar datasets to find coordinated regulatory elements are extended to more cell types, we will be able to find these types of non-coding drivers, which are currently under-studied due to the difficulty in interpreting the non-coding genome, in other cancers. We believe there should be a significant shift in focus to identify non-coding drivers in all cancer types, and this study describes a powerful methodology to do so. Our findings, that hundreds of cis regulatory regions have driver potential, overall support a model under which cancer tumorigenesis and progression is a complex phenomenon, driven by hundreds of driver mutations with different effect sizes, coming both from the somatic and the germ-line genome both coding and non-coding, play an important role in the process. Under this model the well-known protein-coding somatic driver mutations in key genes are the necessary components for tumorigenesis and define many of the common properties of tumours, while the larger number of rarer driver mutations define the specific characteristics of each tumour. It is likely that the specific progression characteristics as well as the response of such tumours to specific therapies and our immune system depend on these larger sets of rare mutations in much the same way they do in complex diseases and this hypothesis provides a strong rationale to explore the non-coding cancer genome more deeply.

## AUTHOR CONTRIBUTIONS

H.O and E.T.D designed the study. H.O. analysed the data and H.O and E.T.D interpreted the results. O.D. analysed and shared the SGX data, M.W.S and C.H. were involved in data analysis. H.O. wrote the manuscript and E.T.D edited it.

## ACKNOWLEDGMENTS

This research was supported by grants from NIH-NIHM (GTEx), European Commission FP7, European Research Council, and Swiss National Science Foundation. The computations were performed at the Vital-IT Centre for high performance computing of the SIB Swiss Institute of Bioinformatics. We thank all members of the Sysgenetix for their crucial input in data generation and analyses.

